# Floral temperatures of the southern-most flowering *Amorphophallus titanum* and seedling growth following self-pollination

**DOI:** 10.1101/2025.07.15.665007

**Authors:** Janice M. Lord, Kat J. Lord

## Abstract

*Amorphophallus titanum* (Becc.) Becc. (Araceae) is native to the rainforests of Sumatra, and famously produces a single, massive, thermogenic and highly odoriferous inflorescence. Despite having both male and female flowers in close proximity, *A. titanum* does not readily self-pollinate as it is strictly dichogamous, with female flowers receptive on the first night of flowering and pollen released on the second night. We report on external and internal temperatures reached by the most southerly plant in cultivation which flowered in February 2018 in Dunedin, New Zealand. We also report on successful self-pollination of the same plant in January 2021 with pollen collected in 2018 that had been stored at -80^°^C. External spathe and internal spadix temperatures exceeded ambient temperatures on both flowering nights in 2018. Self-pollination in 2021 resulted in copious fruits that matured in 8 months and showed 75% germination success. More than a fifth of all seedings to germinate were achlorophyllous and died within four months, which is consistent with inbreeding depression. Seedling height, leaf production, length of active growth period and corm size varied markedly among the surviving chlorophyllous seedlings, with some producing two leaves in their first year. While we would not recommend the use of self-pollination for propagation of this species due to the likely presence of inbreeding depression, our study does prove that pollen can retain its viability at -80^°^C for at least several years which is an important step towards a pedigree-informed breeding programme among plants in cultivation.

## Introduction

*Amorphophallus titanum* (Becc.) Becc. (Araceae), commonly known as the Titan arum or giant corpse flower, is a rare and remarkable plant native to the rainforests of Sumatra, Indonesia, with an IUCN classification of endangered (Yuzammi and Hadiah 2018; Yusniwati et al. 2024; Royal Botanic Gardens, Kew 2025). It is famous for its enormous, foul-smelling reproductive structure and also produces the largest corm of any angiosperm. In non-flowering years a single, highly-dissected leaf supported by a stout petiole arises from the corm and can reach more than 4 m in height. In a flowering year the corm produces a single spadix surrounded by a modified leaf bract called a spathe, as is typical of the Araceae family. The spathe can reach more than 3 metres in height, making it one of the largest unbranched inflorescences in the world. *A. titanium* is monoecious, with female flowers in a zone at the base of the spadix and male flowers above (Yusniwati et al. 2024; Royal Botanic Gardens, Kew 2025).

*A. titanum* inflorescences emit a strong odor reminiscent of rotting flesh due to the production of a range of volatiles including dimethyl disulfide and dimethyl trisulfide (Liu et al. 2023; Yusniwati et al. 2024). The inflorescence is also thermogenic, meaning that during flowering it produces heat endogenously, which is thought to assist in the dissemination of the pungent volatiles (Claudel 2021; Liu et al. 2023; Yusniwati et al. 2024). While it is known to attract a range of insects in its natural habitat, pollination appears to be most commonly achieved by copro-necrophagous Coleoptera and Diptera (Claudel 2021). Despite having both male and female flowers in close proximity, *A. titanum* does not readily self-pollinate as it is strictly dichogamous, specifically protogynous; the female flowers become receptive to pollen on the first night of flowering, but the male flowers do not release pollen until the following night, by which time the female flowers are thought to be no longer receptive (Claudel 2021). The infructescence of *A. titanium* is comprised of numerous, one or two-seeded berries with a fleshy orange pericarp and hard endocarp and testa (Latifah et al. 2015). Recent examination of two populations in west Sumatra suggests that successful fertilisation and fruit production is uncommon in the wild, however seedlings have been observed (Yusniwati et al. 2024). Seed germination is generally slow and likely inhibited by the fruit wall and testa (Latifah et al. 2015). While fruits have long been thought to be dispersed by hornbills, recent observations of wild populations suggest this may not be the case (Yusniwati et al. 2024).

*A. titanum* has been cultivated in botanic gardens throughout the world for more than 140 years and early demand for corms may have contributed to the decline of wild populations (Hetterscheid and Ittenbach 1996). Cultivated plants now, however, may be important for the continued survival of the species (Murrell et al. 2025). The oldest plant in cultivation, in the Royal Botanic Gardens, Kew, first flowered in 1889 (Royal Botanic Gardens, Kew 2025). Hand cross-pollination of plants in cultivation has been attempted in many countries, however relatively few attempts have resulted in successful seed production in cultivation (Sudermono et al. 2016), most likely because two plants need to be flowering slightly asynchronously. Understanding pollen storage options is therefore important for successful cross-pollination. Lobin et al. (2007) reported that *A. titanum* pollen can be stored in liquid nitrogen, or kept at 5^°^C for up to six weeks, with no difference in seed set between stored vs freshly collected pollen when subsequently used. Sudarmono et al. (2016) successfully cross-pollinated a cultivated plant of *A. titanium* in the Bogor Botanic Gardens, Indonesia, with pollen collected from a different plant that had flowered 3 months prior. The pollen had been stored at 5^°^C and still showed high viability, resulting in c.50% seed set. In 2016 Chicago Botanic Garden successfully cross-pollinated a plant with pollen from two donor plants. They found considerable differences in fruit size between the two pollen donors and attributed this to inbreeding depression as the pollen recipient and one of the pollen donors were siblings (Zaworski 2016). Recently, a global pedigree analysis by Murrell et al. (2025) reviewed relatedness among cultivated plants and found that 27% of the 45 crosses documented in their study were between related individuals. They suggested that both pollen banking and pedigree-informed exchanges were needed to preserve genetic diversity among cultivated *A. titanum*.

### *Amorphophallus* in New Zealand

The original specimens of *A. titanum* held by Auckland Domain Wintergarden, Christchurch Botanic Gardens and Dunedin Botanic Garden very likely came from the same source. The Dunedin plant, DBG 20081359, was donated as a corm in 2008 by a private Auckland collector, who had propagated it from seed imported from the Netherlands (*pers. comm.* Tom Myers). Three corms were also donated to Christchurch Botanic Gardens around the same time very likely by the same collector and clones have since been propagated from natural corm divisions (*pers. comm.* Darren Tillet). The original Auckland Domain Wintergarden plants were likely donated by the same collector in 2008 (Cameron 2014) but died during Covid pandemic lockdowns. More have since been propagated there from seeds imported from the U.S.A. (*pers. comm.* Jonathon Corvisy). The plant at Auckland Zoo was acquired in 2019 (*pers. comm.* Ben Goodwin to Hugo Baynes). It was likely propagated clonally from a plant in Austen’s Exotic Garden, Northland (*pers. comm*. Toni Austen). None of the New Zealand plants were included in the pedigree analysis of Murrell et al. (2025) due to uncertainties around their origin (*pers. comm.* Olivia Murrell).

Here we describe the first flowering of *A. titanium* in the Dunedin Botanic Garden in February 2018, which is the most southerly record of flowering globally (Wikipedia 2025). This plant, DBG 20081359, was donated as a corm to Dunedin Botanic Garden in 2008 by a private Auckland grower, who had propagated it from seed imported from the Netherlands (*pers. comm.* Tom Myers). We report on floral temperatures reached by the spathe and spadix over the course of flowering, and describe methods used to collect pollen in 2018. We also report on the successful self-pollination of the same plant using its own frozen pollen when it flowered for a second time in January 2021, and document the viability of subsequent seeds, as well as seedling growth and survival following this unique self-pollination intervention.

## Methods

A single plant of *A. titanium* in cultivation at the Dunedin Botanic Garden Tropical House, New Zealand, showed signs of flowering in early 2018. The progress of bud development and flowering was recorded using timelapse video (https://www.youtube.com/watch?v=H6P6p9xXEY0). From February 2^nd^ just prior to the spathe opening, until the end of flowering, the external temperature of the spathe and internal temperature of the spadix (when visible) was measured at intervals with a Fluke T120 Thermal Imager (Fig. 1A-D) The spathe opened on the night of 3^rd^ February 2018 (Fig 1B, Fig. 2A) at which time the female flowers at the base of the inflorescence were glistening so were presumed to be receptive (Sudermono et al. 2016). During the following day a c.10 × 20cm window was cut into the base of the spathe with a breadknife and trays made from folded aluminium foil were positioned inside below the male flowers (Fig, 1C, Fig. 2B). The window was resealed by replacing the plug of spathe wall material, which was similar in texture to polystyrene, and covering the joins with plastic cling-film and cellulose tape. During the night of 4^th^ February, the male flowers released copious pollen onto the trays. On the morning of 5^th^ February (Fig. 1D), the captured pollen was scraped into falcon tubes and transferred to a -80^°^C freezer in the Department of Botany, University of Otago, Dunedin. A small sample of the fresh pollen was also incubated in a 5% sugar solution to test viability.

**Fig 1:**
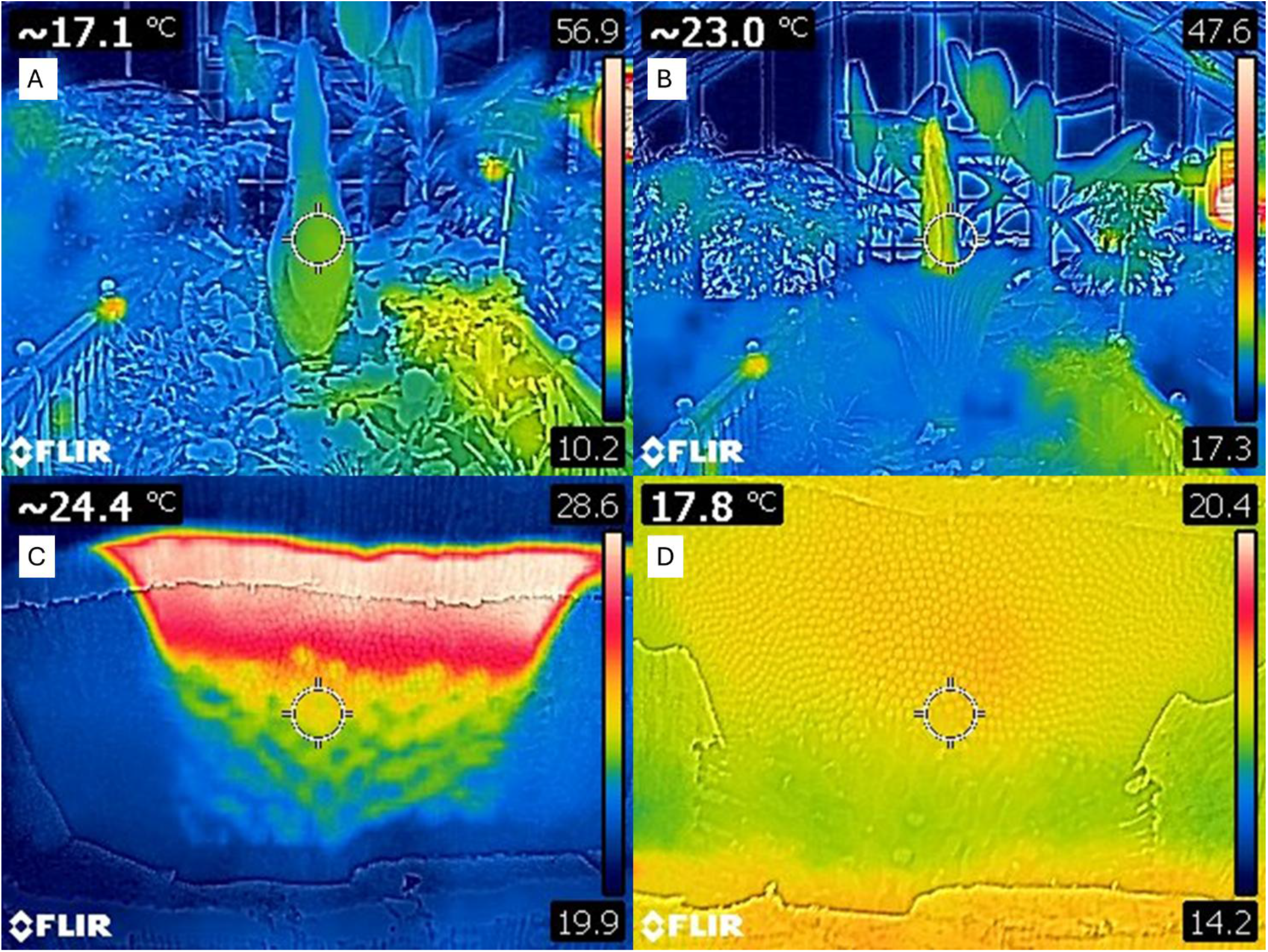
Artificially coloured thermal images of *Amorphophallus titanum* prior to and during flowering in 2018, Dunedin Botanic Gardens. Colour scale differs among panels. Temperature in upper left of each panel corresponds to spot temperature of central target. A: 11.00pm, 2 February, prior to flowering; B: 6.50pm, 3 February, flowering commencing; C: 9pm, 4 February 2018, view through window cut in base, male flowers (red) commencing pollen release, female flowers (green-blue below male flowers) no longer receptive; D: 8.30am, 5 February 2018, view through window cut in base, post-pollen release.

**Fig 2:**
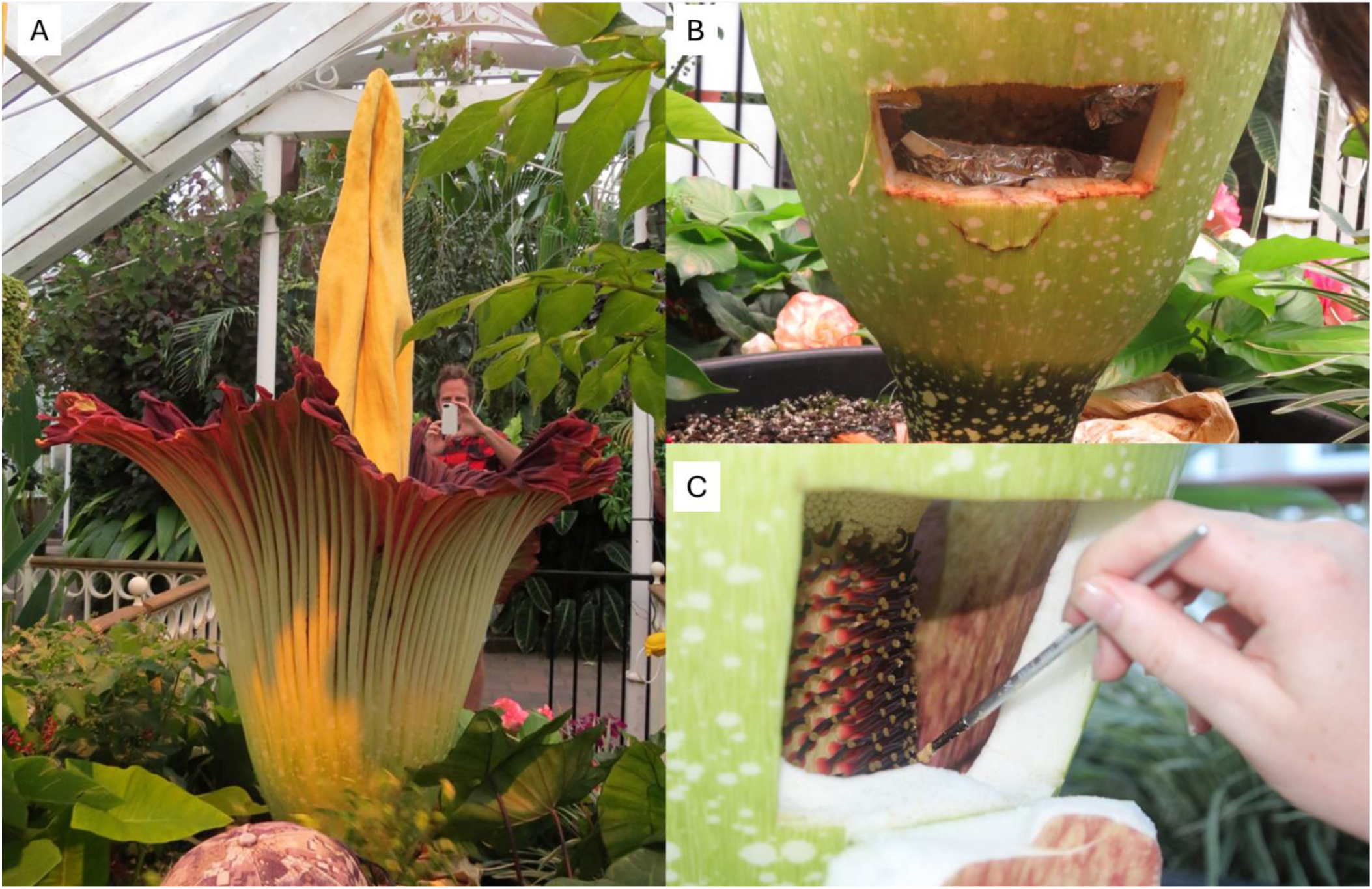
Pollen collection and hand self-pollination of *Amorphophallus titanum,* Dunedin Botanic Gardens. A: Inflorescence after first night flowering, 4 February 2018; B: pollen collected in foil trays following the second night flowering, 5 February 2018; C: pollen that had been stored at -80^°^C being applied to receptive female flowers of the same plant, 15 January 2021 (photos A and B, Tor Sollie; photo C, Meagan Burton).

At the time, in 2018, the intention was to use the stored pollen to cross-pollinate the next New Zealand plant to flower, however when plants in Auckland and Christchurch flowered in 2020, we missed the first night of flowering when female flowers were receptive. When the same Dunedin plant from which pollen was collected flowered again in January 2021, we unable to locate pollen from another plant in time for the first night of flowering in Dunedin so we decided to attempt self-pollination. The spathe of the Dunedin plant began opening on 15^th^ January 2021. In the early evening of that same day when the spathe had opened fully, a hole was cut in the base of the spathe as in 2018 and thawed 2018 pollen was applied freely to stigmas using a paint brush (Fig. 2C). On the second night of flowering (16^th^ January 2021) pollen was collected in aluminium trays as before. Half of this pollen was transferred immediately to the University of Otago, Botany Department -80^°^C freezer, and the other half was stored dry at 4^°^C for 24 hours before also being frozen at -80^°^C. A small sample of the fresh pollen was also incubated in a 5% sugar solution to test viability as in 2018.

Fruit development was monitored and berries collected on 13^th^ September 2021. The following day whole fruits were sown with the fleshy pericarp intact into trays of seed raising mix (¼ perlite, ¼ horticultural sand, ¼ fine bark, ¼ (peat, gravel, sand mix), and 7-8 month osmocote fertiliser) in a heated greenhouse (22^°^C day, 18 ^°^C night, 60% RH). Seedlings were pricked out as they emerged and replanted into individual SP90 (90mm diameter) pots. At approximately four months of age seedlings were repotted into PP150 pots (1.5L).

A subset of 23 seedlings were selected for ongoing growth measurements. Seedling heights were measured at fortnightly intervals over the summers of 2021-2022 and 2022-2023. Height was measured by straightening the leaf lobes to their maximum length (Fig 3). In winter (June) 2024 all 23 pots were emptied so that corms could be inspected and measured. The presence of an intact corm was taken as indicating dormancy rather than death. Pearson’s correlations and one-way ANOVA were used to explore relationships among maximum seedling height prior to winter, length of growth period before dormancy, whether a seedling produced one or two leaves in its first growing season and was still actively growing after 15 months, and corm size at the end of the study. All analyses were conducted in SPSS v.29 (IBM Corporation). Continuous variables were tested for assumptions of normality and homogeneity of variance prior to analyses.

**Fig 3:**
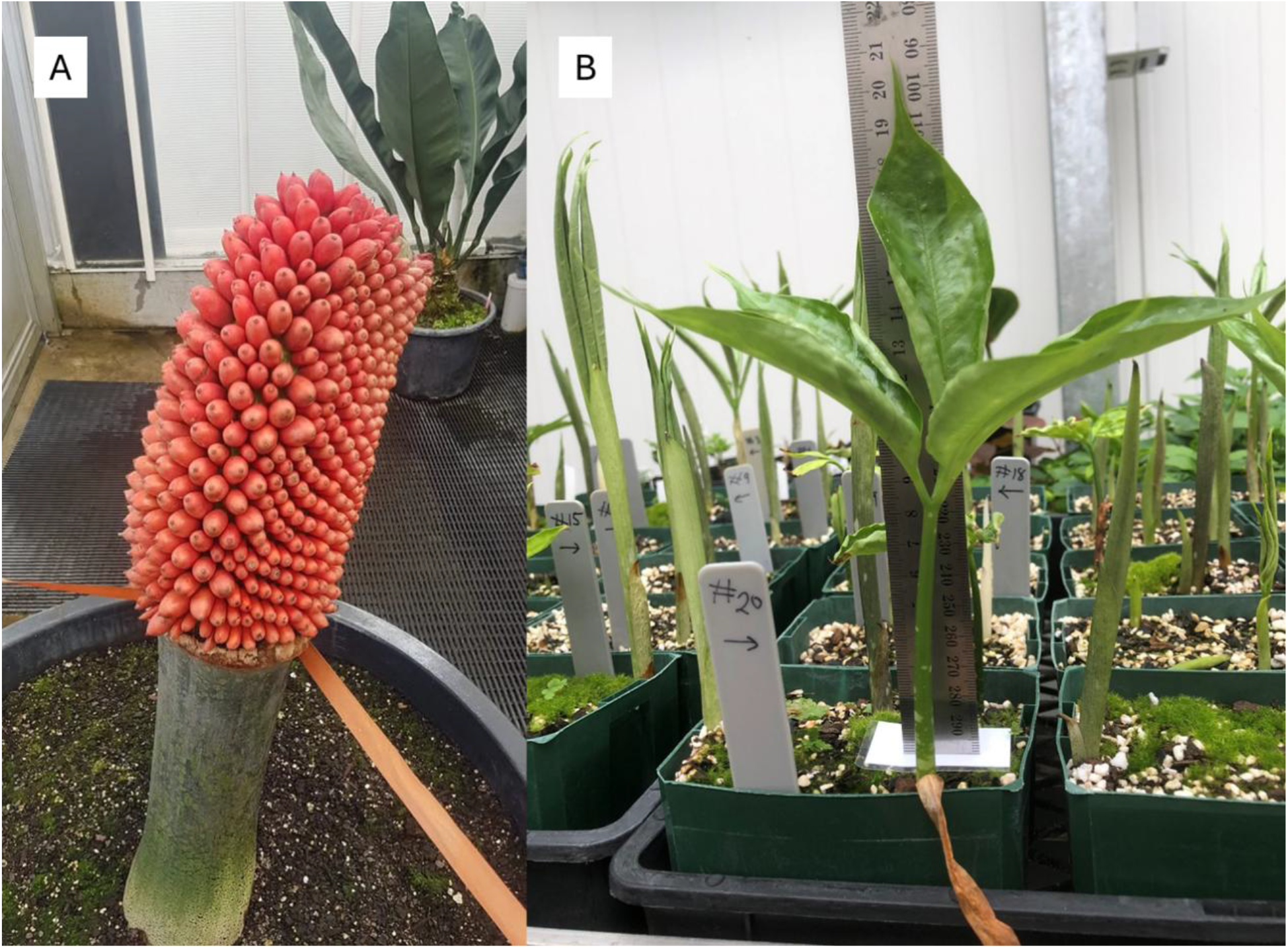
Outcome of *Amorphophallus titanum* self-pollination: (A) mature infructescence, (B) 2-month-old seedlings. (A, Stephen Bishop; B, Kat Lord).

## Results

Thermal imaging captured significant temperature increases externally and internally as flowering progressed in 2018 (Fig. 1A-D). The central spathe was noticeably warmer than ambient temperature of 17^°^C at the start of the first night of flowering, exceeding 23^°^C (Fig. 1B). During this first night the female flowers were presumed to be receptive, and the odour was strongest. On the second night of flowering, during which male flowers dehisced, the internal temperature in the zone of male flowers exceeded 28^°^C, considerably warmer than the zone of female flowers (Fig. 1C). The morning after the second night of flowering the interior of the inflorescence was again at more-or-less ambient temperature (Fig. 1D). The inflorescence proceeded to senesce over the following days despite a few fruits apparently starting to develop.

Pollen freshly collected in 2018 and 2021 readily germinated in 5% sucrose with >50% of grains producing pollen tubes within 48 hours. Following application of stored 2018 self-pollen to female flowers on the first night of flowering in January 2021, fruits were observed developing across the entire infructescence. These were collected for propagation after 8 months when they were bright orange (Fig. 3A). Seedling emergence was first observed at the start of October 2021 approximately 4 weeks after sowing and seedling emergence continued over several days (Fig. 3B). Approximately 75% of seeds sown germinated, resulting in 178 seedlings (Table 1). A substantial number of seedlings emerging were distinctly lacking in chlorophyll, but their proportion of the total surviving seedling cohort decreased over time (Table 1). The subset of 23 seedling used for ongoing height measurements included three achlorophyllus seedlings which all died within four months. Excavation at the end of the study confirmed these seedlings had not developed corms so were assumed to have died by five months (Fig. 5). All 20 chlorophyllous seedlings continued to actively grow for at least six months, however only six still had leaves after 15 months.

**Table 1:** Survival of achlorophyllous (“white”) and chlorophyllous (“green”) seedlings over 18 months post-germination, and average height (mm) of 23 selected seedlings (3 “white”, 20 “green”) over 13 months. The 3 “white” seedlings being measured had died by December 2022.

| *Month* | *total seedlings* | *% white* | *Mean height white (mm)* | *% green* | *Mean height green (mm)* |
| --- | --- | --- | --- | --- | --- |
| Dec 2021 | 178 | 23.0 | 119.67 | 77.0 | 192.00 |
| Jan 2022 | 169 | 21.9 | 237.67 | 78.1 | 309.70 |
| Feb 2022 | 157 | 18.5 | 231.33 | 81.5 | 365.55 |
| Dec 2022 | 151 | 0 | 0 | 100 | 266.35 |
| May 2023 | 132 | 0 | - | 100 | - |

For chlorophyllous seedlings, height growth was relatively rapid with the largest seedlings exceeding 200 mm within 2 months (Fig. 3) and continuing to grow rapidly for the season, some exceeding 600 mm in extended leaf length by the following winter (June 2022), 8 months after germination. Many seedlings retained leaves over their first winter (2022) rather than undergoing a period of dormancy, and more than 50% had produced a second leaf by that time. The maximum height of seedling leaves prior to winter 2022 was positively correlated with the number of weeks before seedlings became dormant (two-tailed Pearson’s correlation: r = 0.762, P < 0.01, N = 20). Seedlings that had produced a second leaf by winter 2022 also achieved a significantly greater maximum leaf height and grew longer before entering dormancy compared with seedlings that only produced one leaf in their first growing season (one-way ANOVA: max height *F*_1,18_ = 27.918, *P* < 0.001; weeks to first dormancy *F*_1,18_ = 8.062, *P* < 0.05).

Corm size varied dramatically among the 23 study seedlings (Fig. 4) with the largest corm measuring 110 mm diameter but most other corms being considerably smaller. One of the fruits sown (#15) produced two corms (Fig. 4). This plant had also produced two leaves by winter 2022. Several other instances of multiple corms in a pot were also recorded during routine maintenance of the 100+ seedlings not being monitored for height. Corm diameter was not significantly correlated with maximum leaf height or weeks to first dormancy; neither did it differ significantly between seedlings that produced one or two leaves prior to winter 2022.

**Fig 4:**
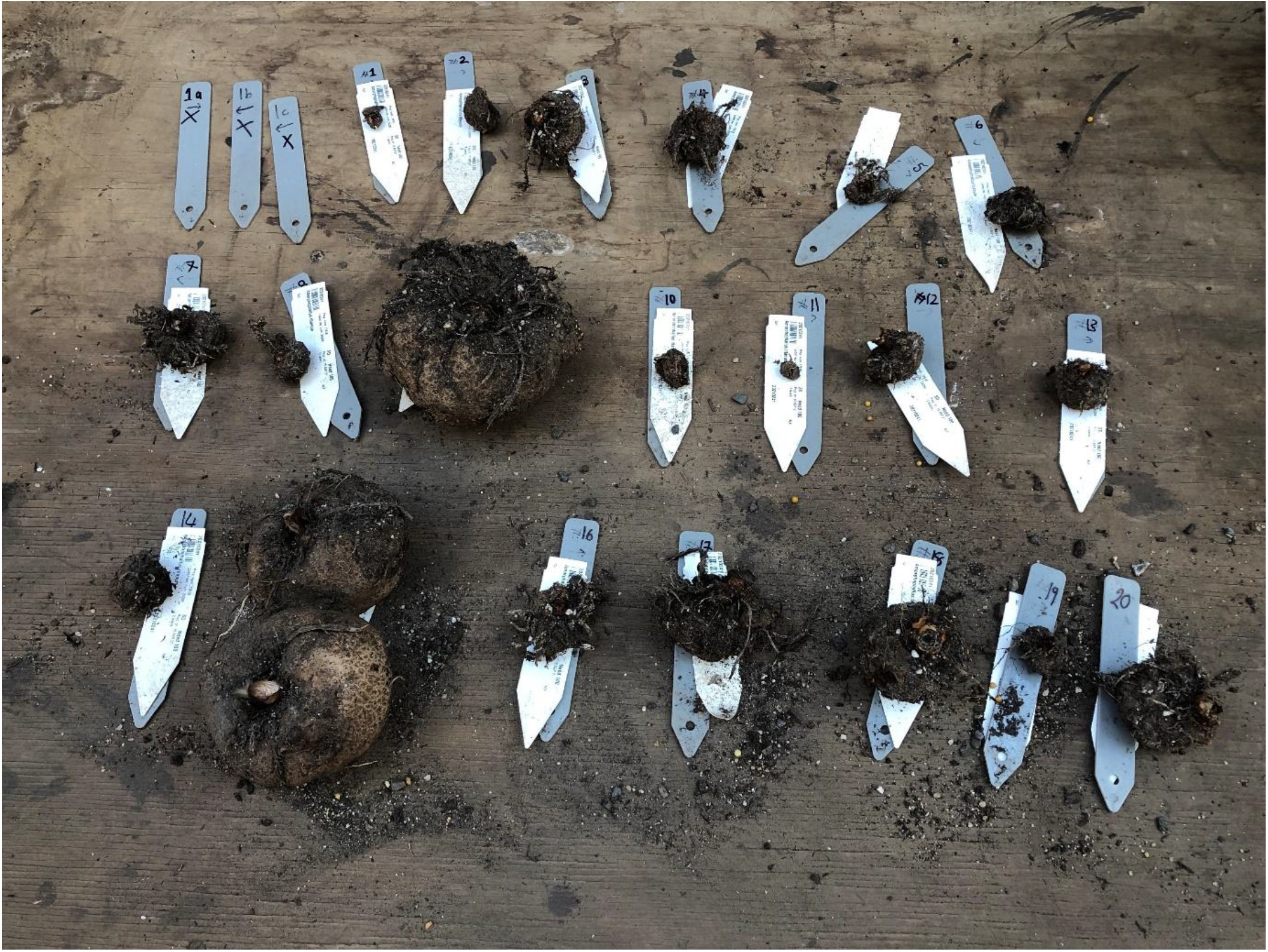
Corms from 23 study seedlings c.32 months after germination. The largest corm, #9, is 110 mm diameter. The three labels marked X in the top left were from pots containing achlorophyllous seedlings in which no corms were found. Note pot #15 (lower left) contained two corms.

## Discussion

This study documented floral temperatures for the most southerly flowering *Amorphophallus titanum* in cultivation and also successfully self-pollinated the same plant. The changes in temperature we recorded as flowering progressed over several nights are consistent with previous findings of thermogenesis for this species, and for the genus in general (Korotkova and Barthlott 2009; Claudel et al. 2023). However, the maximum temperatures we recorded during the evening of flowering nights were somewhat lower than maximum temperatures recorded later during each night (e.g., 36-38^°^C at midnight, Korotkova and Barthlott 2009). Despite this, we were able to record the temperature differential between male and female flowers during the stage at which the inflorescence was switching from pollen receipt /female receptivity to pollen export modes. Our measurement of temperatures leading up to pollen release match previous records that thermogenesis can raise male flower temperatures by more than 10^°^C above ambient (Korotkova and Barthlott 2009).

We were able to self-pollinate the solitary *A. titanum* in Dunedin Botanic Garden using its own pollen that had been stored at -80^°^C for three years. In this manner we circumvented the species’ own barriers to self-pollination and demonstrated that *A. titanum* does not possess a genetic self-incompatibility system. Monoecious species are no more likely to be self-incompatible than hermaphroditic species (Bergin 1993). Instead, spatial (herkogamy) and/or temporal (dichogamy) separation of male and female functions reduces geitenogamous self-pollination among flowers on the same plant (Lloyd and Webb 1986; Webb and Lloyd 1986). *Amorphophallus titanum* demonstrates monoecious interfloral herkogamy and dichogamy in its inflorescence structure; female flowers are located below the zone of male flowers on the central spathe and are receptive before the male flowers release pollen. The observation of a small number of fruits on the plant in Dunedin Botanic Garden in 2018 suggests this species is capable of self-pollination to a limited extent unaided. However, the presence of achlorophyllous seedlings and the wide range of corm sizes among the cohort of chlorophyllous seedlings we managed to raise following forced self-pollination in 2021 suggest the presence of inbreeding depression, as suggested by Zaworski (2016).

Although our seedlings derived from self-pollination they do provide some insights into seedling growth and resource allocation. Seedling leaves were large and grew rapidly, which may relate to the natural shaded forest environment of this species (Yusnawati et al. 2024) in which large seeds and large seedlings would likely have an advantage during establishment (Westoby et al. 1996). Interestingly, seedling height was not correlated with corm size suggesting resource allocation to above and below ground organs is not linked. It was also surprising that more than 50% of seedlings produced two leaves in the first growing season, as adult plants only produce a single leaf annually when in the vegetative stage (Yusnawati et al. 2024, Royal Botanic Gardens, Kew 2025). The finding of multiple corms in some pots is undoubtedly due to some fruits containing two seeds instead of one. As we did not excise seeds from fruits before sowing, we cannot confirm the frequency of two-seeded fruits following self-pollination. However, wild plants are known to produce fruits with one or two seeds (Yusnawati et al. 2024) and two-seeded fruits have been observed following cross-pollination of plants in cultivation (Zaworski 2016).

In light of the risk of inbreeding depression we would not recommend the routine use of forced self-pollination for propagation of isolated plants and also advise against pollination between closely related plants. We support the recommendations of Murrell et al. (2025) that institutions with this species in cultivation share pedigree data so that cross-pollinations can occur between unrelated plants. However, a pedigree-based management approach for *A. titanum* also requires that pollen can be stored, as cultivated plants seldom flower synchronously. In addition to reports of successful short-term pollen storage (e.g. Lobin et al. 2007, Sudarmono et al. 2016, United States Botanic Garden 2023), our study demonstrates that *A. titanum* pollen can remain viable for at least three years at -80^°^C and be used for successful pollination, despite having a trinucleate pollen type which is thought to have reduced longevity. Knowledge of pollen storage potential can provide critical support for the conservation of rare plants (van der Walt et al. 2022, Wolkis et al. 2024) and avoid issues with inbreeding among cultivated populations, as has been the case for many *A. titanum* crosses (Murrell et al. 2025). Co-ordinated pollen banking and sharing of detailed pedigree information among institutions is needed to protect the genetic diversity of this species and other rare plants in cultivation.

## Acknowledgements

Dunedin Botanic Garden management for after-hours access and permission to work with the plant, DBG staff especially Stephen Bishop, Tom Myers and Alice Lloyd-Fit for plant expertise and project support, Tor Sollie, Jessica Paull, Meagan Burton and Kate Moss-Mason for assisting with pollen collection and pollination. Jonathon Corvisy (Auckland Domain Wintergarden), Hugo Baynes and Ben Goodwin (Auckland Zoo), and Darren Tillet (Christchurch Botanic Gardens) for information on plant origins. Olivia Murrell for additional pedigree information. Botany Department, University of Otago for use of equipment and low temperature storage space and the Brenda Shore Trust for financial support.

## References

Bertin RI. 1993. Incidence of monoecy and dichogamy in relation to self-fertilization in angiosperms. Am. J. Bot. 80: 557–560. 10.1002/j.1537-2197.1993.tb13840.x

Bhatia R. 2024. Rare corpse flower blooming at Auckland Zoo. Stuff online, 4 December. https://www.stuff.co.nz/travel/360509940/rare-corpse-flower-blooming-auckland-zoo.

Cameron EK. 2014. Titan arum (Amorphophallus titanum) flowers in New Zealand for the first time. Auck. Bot. Soc. J. 69:85–88. https://bts.nzpcn.org.nz/site/assets/files/24066/ak_bot_soc_journal_69_1_jun_2014_85-88.pdf

Claudel C. 2021. The many elusive pollinators in the genus Amorphophallus. Arthropod-Plant Inter. 15:833 –844. 10.1007/s11829-021-09865-x

Claudel C, Loiseau O, Silvestro D, Lev-Yadun S, Antonelli A. 2023. Patterns and drivers if heat production in the plant genus Amorphophallus. Plant J. 115:874–894. 10.1111/tpj.16343

Hetterscheid W, Ittenbach S. 1996. Everything you always wanted to know about Amorphophallus, but were afraid to stick your nose into!!!!! J. Int. Aroid Soc. 19:7–131. https://www.researchgate.net/publication/283567478

Korotkova N, Barthlott W. 2009. On the thermogenesis of the Titan arum (Amorphophallus titanum) Plant Signal. Behav. 4: 1096–1098. 10.4161/psb.4.11.9872

Latifah D, Purwantoro RS. 2015. Seed germination of the Giant Corpse Flower Amorphophallus titanum (Becc.) Becc. Ex Arcang: the influence of testa. Berita Biologi 14:39–47. 10.14203/BERITABIOLOGI.V14I1.1861

Liu D, Zhang P, Liu D, Feng Y, Chi M, Guo Z, Wang X, Zhing J, Sun M. 2023. An analysis of volatile compounds and study of release regularity in the flower of Amorphophallus titanum in four periods. Horticulturae 9: 487. 10.3390/horticulturae9040487

Lloyd DG, Webb CJ. 1986. The avoidance of interference between the presentation of pollen and stigmas in angiosperms. I. Dichogamy. New Zeal. J. Bot. 24: 135–162. https://www.tandfonline.com/doi/abs/10.1080/0028825X.1986.10409725

Lobin, W, Neumann M, Radscheit M, Barthlott W. 2007. The Cultivation of Titan Arum (Amorphophallus titanum): a Flagship Species for Botanic Gardens”, Sibbaldia 5: 69–86. doi:10.24823/Sibbaldia.2007.8.

Murrell OG, Diaz-Martin Z, Havens K, Hughes M, Meyer A, Tutt J, Zerega N, Fant JB. 2025. Using pedigree tracking of the ex situ metacollection of Amorphophallus titanum (Araceae) to identify challenges to maintaining genetic diversity in the botanical community. Ann. Bot.: mcaf038, 10.1093/aob/mcaf038

Radio New Zealand. 2020. ‘Corpse flower’ bloom to cause a stink at Auckland Wintergarden. RNZ online, 3 January. https://www.rnz.co.nz/news/national/406638/corpse-flower-bloom-to-cause-a-stink-at-auckland-wintergarden

Royal Botanic Gardens, Kew. 2025. Titan Arum. https://www.kew.org/plants/titan-arum. Accessed 29 January 2025.

Sudarmono, Latifah D., Hartini S., Wawangningrum H. 2016. Hand-pollination of the giant corpse flower in the Bogor Botanic Gardens. Int. J. Conserv. Sci. 7:1153–1160. https://ijcs.ro/public/IJCS-16-64_Sudarmono.pdf

United States Botanic Garden. 2023. 2021 Corpse Flower. https://www.usbg.gov/2021-corpse-flower. Accessed 3 March 2025.

Van Beynan J, Walters L. 2015. Thousands flock to see Auckland Wintergarden’s ‘corpse flower’. Stuff online article June 17. https://www.stuff.co.nz/auckland/69430894/thousands-flock-to-see-auckland-wintergardens-corpse-flower

van der Walt K, Alderton-Moss J, Lehnebach CA. 2023. Cross-pollination and pollen storage to assist conservation of Metrosideros bartlettii (Myrtaceae), a critically endangered tree from Aotearoa New Zealand. Pac. Cons. Biol. 29:141 –152. 10.1071/PC21054

Webb CJ, Lloyd DG 1986. The avoidance of interference between the presentation of pollen and stigmas in angiosperms. II. Herkogamy. New Zeal. J. Bot. 24: 163–178. https://www.tandfonline.com/doi/abs/10.1080/0028825X.1986.10409726

Westoby M, Leishman MR, Lord JM. 1996. Comparative ecology of seed size and dispersal. Phil. Trans. Roy. Soc. Lond. B 351: 1309–1318. 10.1098/rstb.1996.0114

Wikipedia. 2025. List of publicised titan arum blooms in cultivation. https://en.wikipedia.org/wiki/List_of_publicised_titan_arum_blooms_in_cultivation Accessed 19 March 2025.

Wolkis D, Eltringham C, Fant J, Foster J, Knight T, Meyer A, Romero-Saltos H, Walsh SK, Wood A, Havens K. 2024. Pollen banking is a critical need for conserving plant diversity. Nat. Plants 10:1270 –1271. 10.1038/s41477-024-01757-1

Yusniwati, Setiawan RB, Handayani M, Nanda AR, Sukma D, Rahmi A, Syahputra A, Bosma PAL, Baiturrahman A. 2024. Expedition and Characterization of the Corpse Flower (Amorphophallus titanum Becc.) in West Sumatra. J. Manajemen Hutan Tropika 30: 258–264. 10.7226/jtfm.30.2.258

Yuzammi, Hadiah JT. 2018. Amorphophallus titanum. The IUCN Red List of Threatened Species 2018: e.T118042834A118043213. 10.2305/IUCN.UK.2018-2.RLTS.T118042834A118043213.en. Accessed 29 January 2025.

Zaworski K. 2016. Alice the Amorphophallus—An Update on Titan Arum Fruit. Chicago Botanic Garden, Plant Science and Conservation online article. https://www.chicagobotanic.org/blog/plant_science_conservation/alice_amorphophallus_an_update_titan_arum_fruit

